# SAHA increases chaperone expression and reduces Z-alpha-1-antitrypsin polymers in a patient specific iPSC-based liver model for alpha-1-antitrypsin deficiency

**DOI:** 10.64898/2026.03.16.711579

**Authors:** Nina Graffmann, Rabea Hokamp, Christiane Lörch, Malin Fromme, Wasco Wruck, Pavel Strnad, James Adjaye

## Abstract

The most severe phenotype of alpha-1-antitrypsin deficiency (AATD) is caused by the Z-mutation within the *SERPINA1* gene. The Glu342Lys substitution causes misfolding and polymerisation of the alpha-1-antitrypsin (AAT) protein, its accumulation in the ER and increases the susceptibility of hepatocytes towards ER-stress. Here, we present an induced pluripotent stem cell (iPSC)-based hepatic model to study AATD. We demonstrate that iPSCs from AATD patients differentiate equally well to hepatocyte-like cells (HLCs) as control iPSCs. We detected ZAAT polymers in patient-derived HLCs which could be reduced by SAHA or CBZ treatment. Transcriptome analyses revealed major differences in metabolism and signalling between control and AATD HLCs and indicated increased stress levels affecting intracellular organelles. Importantly, the transcriptomes of control and patient-derived cells separated into distinct clusters with respect to the expression of Heat-shock protein (HSP) encoding genes. SAHA treatment increased expression of various HSPs which might contribute towards reduced ZAAT polymers.

## Introduction

Severe alpha-1-antitrypsin deficiency (AATD) is one of the most common genetic disorders worldwide. It is an autosomal co-dominant inherited condition with an estimated prevalence of approximately 1:2,500 - 1:5,000 (Hamesch et al., 2019). The disease is characterised by low serum levels of alpha-1-antitrypsin (AAT) and is associated with an increased risk of the development of lung and/or liver diseases, including lung emphysema, chronic obstructive pulmonary disease (COPD), liver cirrhosis and hepatocellular carcinoma (HCC) (Greulich et al., 2017; Strnad et al., 2020).

AAT is a serine protease inhibitor predominantly synthesized and secreted by hepatocytes (Janciauskiene and Welte, 2016). Its main target is the serine protease neutrophile elastase (NE) which is released by neutrophile granulocytes during inflammation or infection (Ogushi et al., 1987). The high proteolytic activity of NE is primarily directed against bacteria and other pathogens but it is also associated with tissue damage (Voynow and Shinbashi, 2021). AAT is essential to counteract this detrimental effect of NE, especially in the lungs.

The 52 kDa glycoprotein AAT is encoded within the *SERPINA1* gene located on chromosome 14q32 (Seixas and Marques, 2021). *SERPINA1* is a highly polymorphic gene with more than 500 single nucleotide polymorphisms (SNP) being observed to date (Seixas and Marques, 2021). These polymorphisms can lead to changes in the general AAT expression efficiency as well as modifications of the protein structure itself. The AAT wild-type (WT) form is designated as protease inhibitor (PI)*M (Crystal, 1990). The most common severe *SERPINA1* gene variant which is related to AATD is referred to as Pi*Z. The homozygous Pi*ZZ genotype is seen in approximately 96 % of severe AATD patients that are highly susceptible to clinically manifested liver disease (Blanco et al., 2017). The Z-mutation, which is associated with an 85 % reduction of AAT serum levels, is a single point mutation within the *SERPINA1* gene, leading to the substitution of glutamate at position 342 (Glu342) with lysin (Lys) (Lomas et al., 1992). This fosters misfolding and aggregation of ZAAT, increasing the cells’ susceptibility to endoplasmic reticulum (ER) stress (Ordonez et al., 2013). In line with enhanced ER stress, human primary hepatocytes with ZAAT aggregates as well as IB3 cells overexpressing ZAAT have higher levels of the key ER chaperone 78 kDa glucose-regulated protein (GRP78) (Spivak et al., 2025). ER stress can activate the unfolded protein response (UPR), inducing autophagic and proteasomal degradation of misfolded proteins to protect the protein homeostasis in the cell (Karatas and Bouchecareilh, 2020). If these degradative pathways fail to ensure proteostasis, ER stress-induced apoptosis eliminates the damaged cells (Greene and McElvaney, 2010). Nevertheless, high ER stress levels are associated with the formation of cirrhosis and HCC (Jackson et al., 2023).

Several compounds such as Carbamezepine (CBZ), Suberoylanilide hydroxamic acid (SAHA), Kifunensine (KIF), and Cysteamine (CYS) have shown effects on ZAAT folding *in vitro* or in mouse models (Barthet et al., 2021; Karatas et al., 2021; Marcus and Perlmutter, 2000; Wilson et al., 2015). However, it is difficult to translate these results into the clinic, mainly because of major model immanent drawbacks. Mice express their own complex pattern of *SERPINs* that need to be deleted before a lung phenotype becomes evident, while the liver phenotype can only be triggered by overexpression of the human ZAAT (Borel et al., 2018; Carlson et al., 1989; Khodayari et al., 2021). Likewise, human immortalised cell lines require genetic modifications to allow overexpression of AATD-associated ZAAT (Bouchecareilh et al., 2012). Despite of their human origin, these tumour-derived, modified cell lines often exhibit highly abnormal expression profiles which impair the transferability to the human physiology (Chalak et al., 2024). To overcome these limitations, we and others propose induced pluripotent stem cell (iPSC) derived hepatocyte-like cells (HLCs) as a more natural and effective alternative for liver disease models (Choi et al., 2013; Kaserman et al., 2020; Kaserman et al., 2022; Kaserman and Wilson, 2018; Rashid et al., 2010; Segeritz et al., 2018; Tafaleng et al., 2015; Werder et al., 2021; Wilson *et al*., 2015; Yusa et al., 2011).

In this study, we differentiated iPSCs derived from AATD patients and a control donor into hepatocyte-like cells (HLCs) to study the effect of the above-mentioned small molecules on AAT aggregate formation. We could show that all lines were capable of differentiating into HLCs with the AATD-derived HLCs storing significant amounts of ZAAT polymers which could be reduced by treatment with CBZ and SAHA, the latter inducing the expression of numerous members of the chaperone family.

## Results

### AATD patient-derived iPSCs differentiate efficiently into hepatocyte-like cells (HLCs) and display ZAAT polymers

In order to test if AATD-patient-derived iPSCs can be differentiated into HLCs, we applied our recently optimized protocol (Loerch et al., 2024) to induced pluripotent stem cells (iPSCs) derived from two adult AATD-patients with homozygous Z-mutation (UKA4, 6) (Ncube et al., 2023) and a control donor (UM51) (Bohndorf et al., 2017) (Table 1).

**Table. 1:**
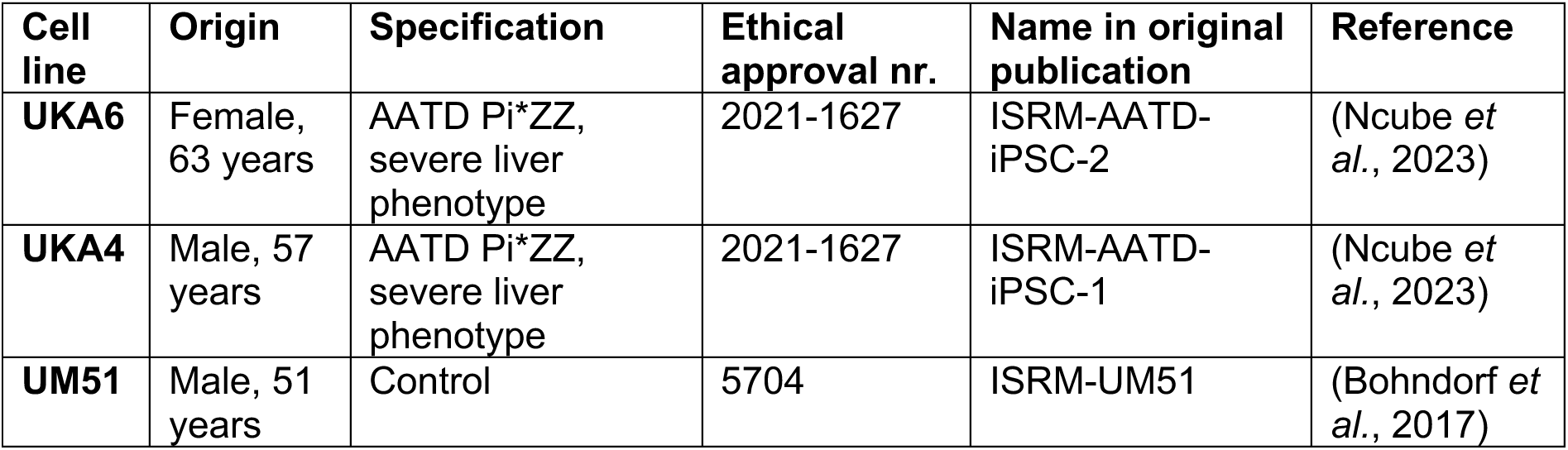
iPSC lines used in this study.

**Tabl. 1: iPSC lines used in this study**
| Cell line | Origin | Specification | Ethical approval nr. | Name in original publication | Reference |
| --- | --- | --- | --- | --- | --- |
| **UKA6** | Female, 63 years | AATD Pi*ZZ, severe liver phenotype | 2021-1627 | ISRM-AATD-iPSC-2 | (Ncube *et al.*, 2023) |
| **UKA4** | Male, 57 years | AATD Pi*ZZ, severe liver phenotype | 2021-1627 | ISRM-AATD-iPSC-1 | (Ncube *et al.*, 2023) |
| **UM51** | Male, 51 years | Control | 5704 | ISRM-UM51 | (Bohndorf *et al.*, 2017) |

Bright-field imaging shows proper differentiation of all 3 lines, with definitive endoderm (DE) cells showing classical petal-like morphology, that shifted towards the polygonal shape in hepatic endoderm (HE) and finally hepatocyte-like cell (HLC) stage, where some binucleated cells were visible (Fig. 1A).

**Fig. 1:**
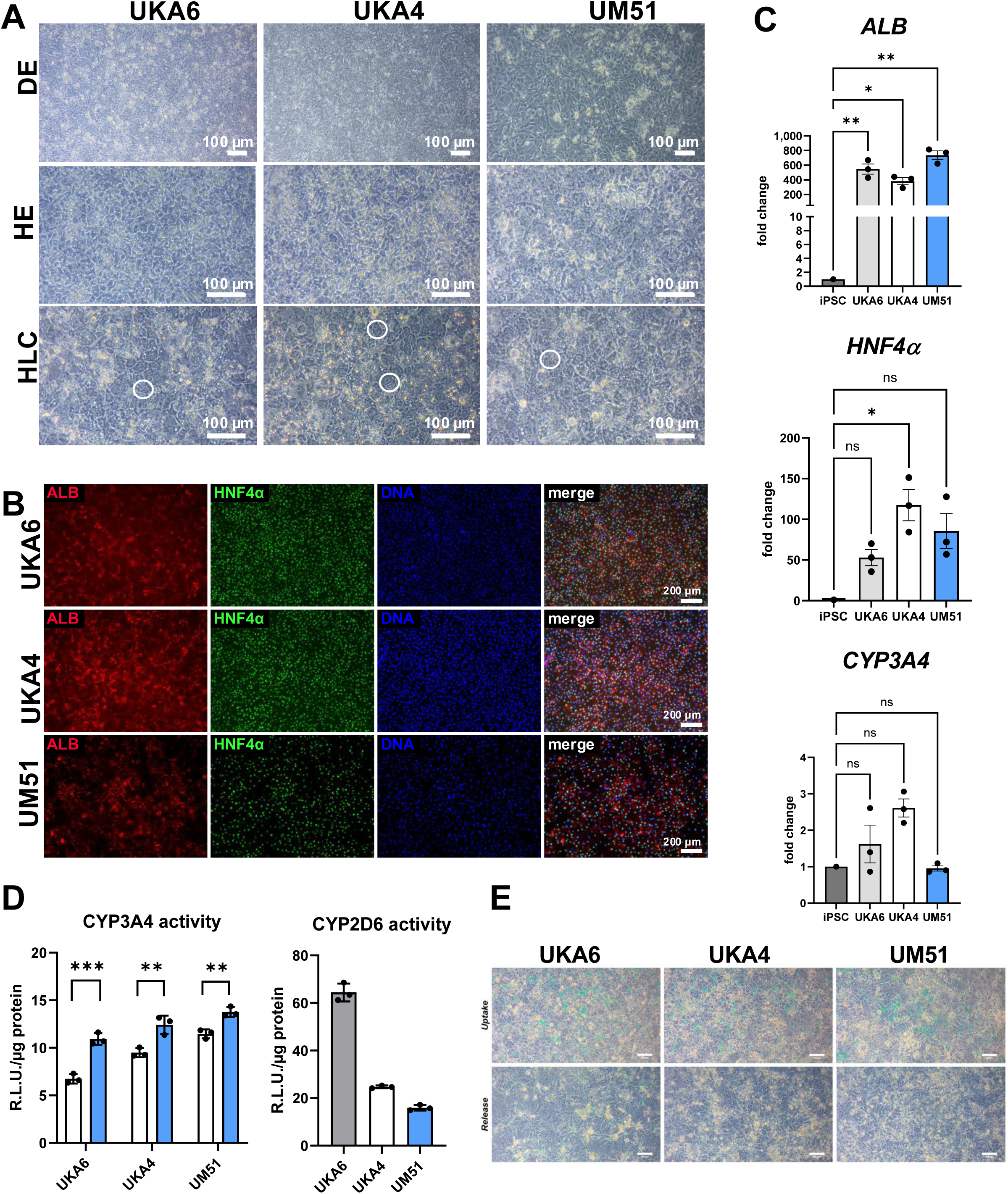
AATD patient-derived iPSCs differentiate with similar efficiency into hepatocyte like cells (HLCs) as cells from a control donor. **(A)** Bright field images show the differentiation process. All iPSCs proceed from petal-shaped definitive endoderm (DE). via hexagonal hepatic endoderm (HE) to cobblestone-like HLC stage with some cells presenting two nuclei (white circles). Scale bar = 100 µM. **(B)** HLCs express the hepatocyte markers albumin (ALB) and hepatocyte nuclear factor 4 alpha (HNF4a). Scale bar = 200 µm. **(C)** Relative mRNA expression of *ALB*, *HNF4a*, and *CYP3A4* normalized to the housekeeping gene *RPLP0*. Bar plots show mean of triplicates for each donor ±SEM. Two-tailed Student’s t-test was performed to calculate significances (*p < 0.05, **p < 0.01, ***p < 0.001). **(D)** CYP3A4 with and without rifampicin (Rifa) treatment for 24 h and CYP2D6 activity were determined by metabolization of luciferin-IPA or luciferin-ME EGE, respectively. Relative light units (R.L.U.) were normalized to total protein. Bar plots show mean of technical triplicates ±SD and significances were calculated by two-tailed Student’s t-test (*p < 0.05, ***p <0.001). **(E)** Uptake of indocyanine green (ICG) and its release after 6 h.

Representative immunocytochemistry shows the expression of Hepatocyte nuclear factor 4 alpha (HNF4α) and Albumin (ALB) in HLC stage (Fig. 1B). QRT-PCR confirmed the expression of *ALB* and *HNF4α* and showed expression of the functional marker Cytochrome P450 family member *CYP3A4* (Fig. 1C). HLC functionality was confirmed for all 3 lines by measuring CYP3A4 and CYP2D6 activity (Fig. 1D) as well as indocyanine green dye uptake and release (Fig. 1E).

Next, we confirmed by qRT-PCR, that all cells expressed *SERPINA1* to a comparable level (Fig. 2A). In Western Blots, we detected lower levels of AAT in controls, while ALB levels were comparable in all lines (Fig. 2B) which is consistent with prior observations from Wilson et al (Wilson *et al*., 2015). Immunocytochemistry showed that all lines expressed AAT, while polymeric ZAAT was only detectable in patient-derived cells (Fig. 2C). Importantly, we observed clump-like staining for AAT and ZAAT in patient-derived cells while control cells showed a more even distribution of AAT in the cytoplasm.

**Fig. 2:**
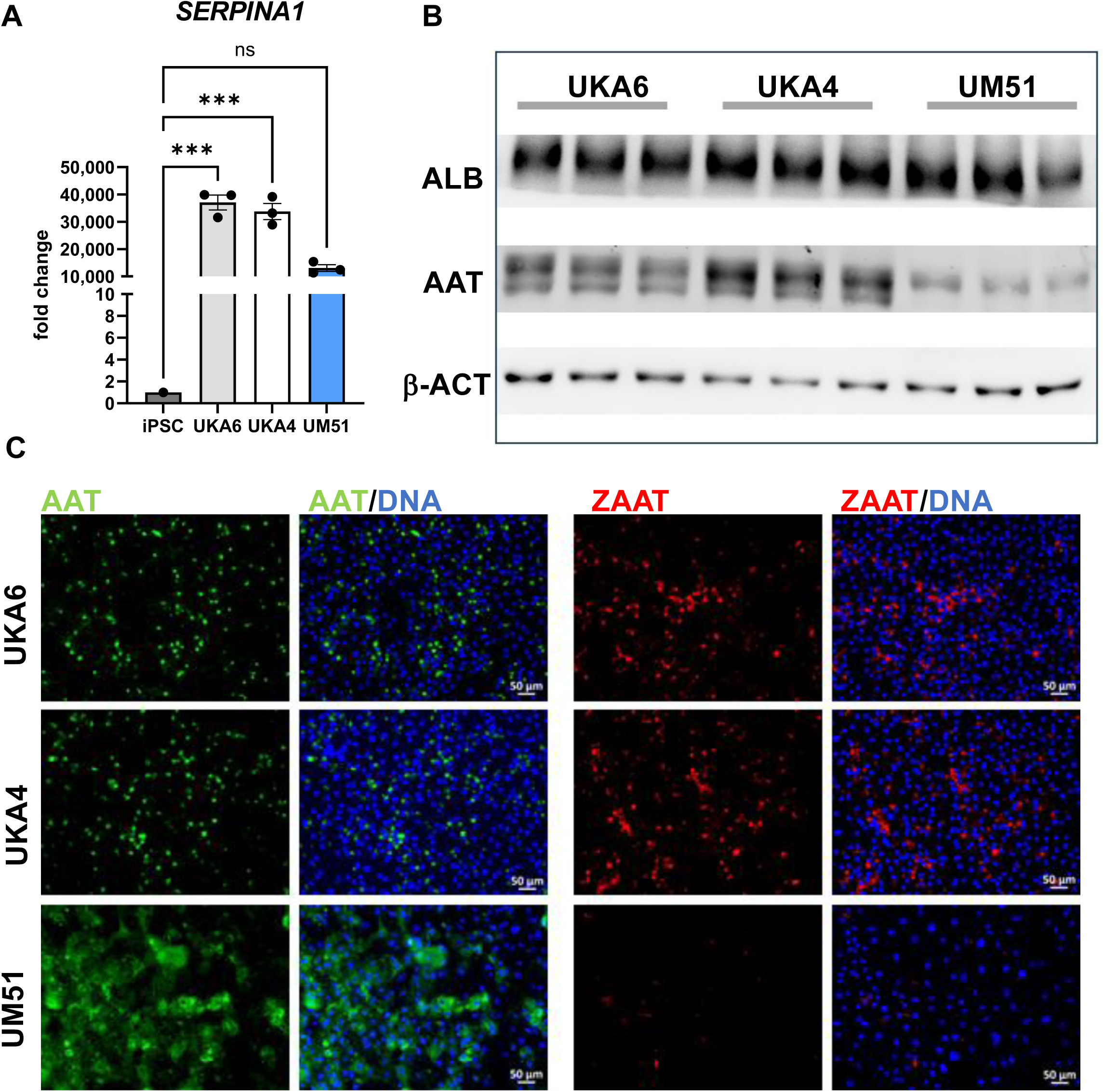
Alpha-1-antitrypsin (AAT) and ZAAT expression in AATD patient-derived and control HLCs. **(A)** All three cell lines express similar levels of *SERPINA1.* Expression was normalized to the housekeeping gene *RPLP0*. Bar plots show mean of biological triplicates ±SEM. Two-tailed Student’s t-test was performed to calculate significances (*p < 0.05, **p < 0.01, ***p < 0.001). **(B)** The hepatocyte marker ALB as well as AAT can be detected in HLCs of all three donors. **(C)** AAT (green) could be detected in all HLCs via ICC. while polymeric ZAAT (red) was only present in AATD patient-derived HLCs. Scalebar = 50 µm

### AATD patient-derived HLCs show dysregulation of heat-shock proteins (HSPs) compared to healthy controls

We performed bulk RNA-seq to analyse the global transcriptomic differences in patient-derived and control HLCs. We identified 14,049 genes expressed in common in both AATD and control donor derived HLCs while 1,082 and 687 genes were exclusively expressed in the control and diseased HLCs, respectively (Fig. 3A). Combining the exclusively and commonly expressed genes, we found 2,052 genes significantly upregulated and 2,617 genes significantly downregulated in patient-derived HLCs (Suppl. Table S4). As ZAAT polymers influence cellular compartments (CC) (Segeritz *et al*., 2018), we checked the top differentially affected Gene-Ontology (GO) CC-terms (p-value < 9 x 10^-12). In line with the data from Segeritz et al. (Segeritz *et al*., 2018), our results show that terms associated with organelles and intracellular membrane enclosed lumens were significantly up-regulated in AATD-derived HLCs compared to the control (Fig. 3B, Suppl. Table S4). Applying a p-value <0.05, we looked for CC-terms specifically related to mitochondria, endoplasmic reticulum and golgi and found these enriched in the up-regulated GO-CC terms in AATD-derived HLCs (Fig. 3C, Supple Table S4).

**Fig. 3:**
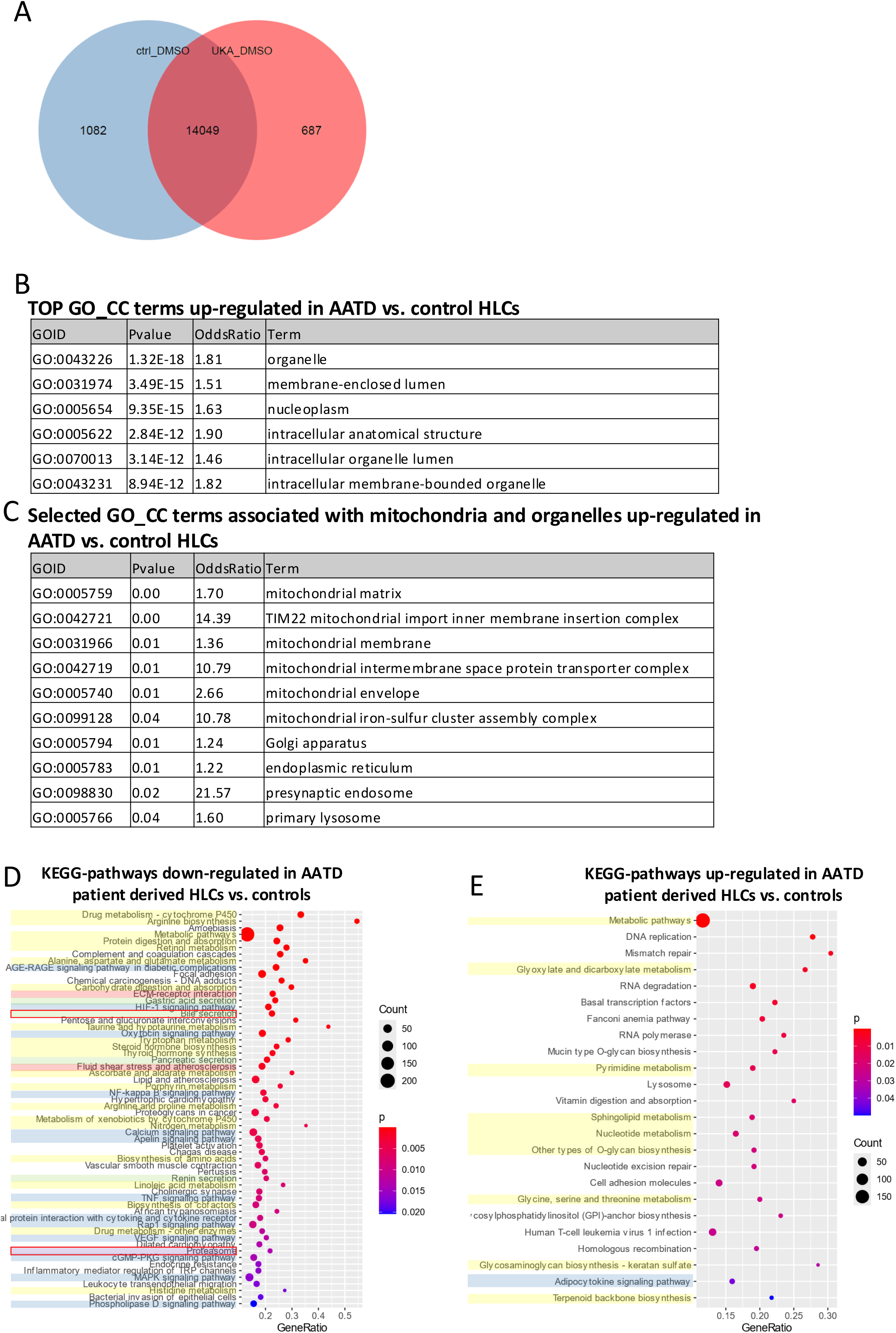
AATD patient-derived HLCs show up-regulated expression of organelle/mitochondria related genes as well as dysregulation of metabolic pathways. **(A)** Venn diagram indicating number of genes exclusively expressed in HLCs derived from a control donor (1,082) and from patient-derived HLCs (687), and commonly expressed genes (14,049) **(B)** GO_CC Terms significantly up-regulated in AATD patient-derived HLCs in comparison to controls. **(C)** GO_CC terms associated with mitochondria and organelles up-regulated in AATD patient-derived HLCs vs. controls. **(D)** KEGG pathways significantly down-regulated in AATD patient-derived HLCs compared to controls. **(E)** KEGG pathways significantly up-regulated in AATD patient-derived HLCs compared to controls. For full data set, please refer to Suppl. Table S4.

A detailed analysis of the affected KEGG-pathways revealed numerous metabolism-associated pathways being either up- or down-regulated in patient-derived cells (Fig. 3D,E, Supplementary Table S4). Several of the down-regulated pathways were associated with amino acid metabolism (“Arginine Biosynthesis”, “Alanine, Aspartate and Glutamate metabolism”, “Tryptophan metabolism”, “Arginine and proline metabolism”, “Biosynthesis of amino acids”, “Histidine metabolism”) which has been recently demonstrated to be disrupted in AATD patients (Kaserman *et al*., 2022). Interestingly, signalling pathways were almost exclusively down-regulated in patient-derived HLCs, comprising, amongst others MAPK, RAP1, cAMP, and calcium signalling, which have been linked to fibrosis, steatosis, cholestasis, and ER stress, respectively (Agarwal et al., 2025; Dabbagh et al., 2001; Fu et al., 2011; Pontisso et al., 2024). STRING-Network analysis of genes that are involved in at least 2 of the KEGG signalling pathways demonstrates their tight interaction (Suppl. Fig. S1). Excretion pathways in general and bile secretion in particular, were down-regulated in AATD-patient-derived cells supporting the phenotype of cholestasis which is observed in paediatric AATD patients. In addition, the proteasome pathway is also downregulated which might be linked to the delayed procession of polymeric AATD (Fig. 3D). Most importantly, we observed a clear clustering of the transcriptomes of control cells versus patient cells when comparing the expression of genes encoding heat shock proteins (HSPs) (Suppl. Fig. S2).

### SAHA and CBZ reduce ZAAT polymers in AATD patient-derived HLCs

To prove that our patient-derived HLCs are suitable for modelling AATD, it is essential that the amount of ZAAT polymers can be reduced with small molecules. Therefore, we incubated patient-derived HLCs with 4 compounds that have previously shown beneficial effects on the disease phenotype in other models. The autophagy enhancer carbamazepine (CBZ) significantly reduced intracellular ZAAT accumulation in murine hepatocytes up to the reversal of AATD-associated fibrosis in a Pi*Z mouse model (Hidvegi et al., 2010). It is the only drug that was also tested in ZAAT patient-derived HLCs, where it could reduce intracellular AAT levels without influencing its secretion (Choi *et al*., 2013; Wilson *et al*., 2015). By inhibiting α-mannosidase I-mediated trimming of mannose residues, kifunensine (KIF) decreased clearance of misfolded proteins thus enhancing ZAAT folding efficiency and secretion rates in fibroblasts overexpressing ZAAT (Marcus and Perlmutter, 2000). Cysteamine (CYS) reduces the activity of Protein Disulfide Isomerase Family A Member 4 which decreases disulfide bond dependent ZAAT polymerisation in Huh7 and IB3 cell lines overexpressing ZAAT (Karatas *et al*., 2021). The histone deacetylase inhibitor (HDACi) SAHA enhances expression of numerous genes, including chaperones and increases the activity of chaperones by maintaining their acetylated state (Bouchecareilh *et al*., 2012; Wang et al., 2017). SAHA increased ZAAT secretion and activity to a level of 50% from wildtype in IB3 and HC116 cells overexpressing ZAAT (Bouchecareilh *et al*., 2012).

In order to determine suitable concentrations of all 4 compounds for the 48 h treatment in our system, we tested for each compound 9 distinct concentrations ranging between 0.1 μM – 400 μM for CBZ and KIF, 0.05 μM – 100 μM for SAHA, and 10 μM – 1 mM for CYS on UKA6 HLCs. ZAAT load was evaluated by ICC for each condition to assess the appropriate concentration which was defined as the minimal concentration showing a distinct effect on ZAAT load within the treated HLCs. A resazurin assay showed no significant signs of cell death after 48 h of treatment with each compound at all tested concentrations (data not shown). Based on these data, we determined 25 μM CBZ, 10 μM SAHA, 25 µM KIF, and 250 µM CYS as suitable concentrations for the following experiments. All selected concentrations were either previously used in the literature or were very close to published effective concentrations (Bouchecareilh *et al*., 2012; Karatas *et al*., 2021; Reiterer et al., 2010; Wilson *et al*., 2015). Suppl. Fig. S3 shows representative pictures for the cells exposed with lowest, highest as well as selected concentration for each component.

ICC based quantification of the ZAAT load revealed a significant reduction in HLCs derived from both AATD-patients in case of SAHA (77.2% reduction in UKA6 and 64.4% in UKA4) and CBZ (52.2% reduction in UKA6 and 54.3% in UKA4) treatment while KIF reduced ZAAT polymers only in UKA4 cells at the selected concentrations (Fig. 4A,B, Suppl. Fig. S4A,B). In parallel with the ZAAT reduction, we also observed a reduction of AAT in HLCs treated with SAHA (37.9% reduction in UKA6 and 38.6% in UKA4) or CBZ (48.8% reduction in UKA6 and 47.7% in UKA4) (Fig. 4A,C, Suppl. Fig. S4A,C). Western Blot analysis of the Triton-X 100 soluble fraction confirmed a down-regulation of AAT after SAHA treatment but not after CBZ treatment (Fig. 4D). However, due to material constrains, we could not repeat this analysis to obtain statistical significance. In case of the Triton-X insoluble faction, we observed very weak bands, which is in line with the fact that ZAAT aggregates grow and mature over time in hepatocytes (Spivak *et al*., 2025) a process that cannot be fully mirrored in cell culture, because of the shorter incubation time and the well-known functional gap between HLCs and primary hepatocytes (Suleman et al., 2025; Wesley et al., 2022). Interestingly though, we observed a reduction not only in ZAAT but also in b-Actin after SAHA treatment, indicating a more global influence of SAHA on proteins in the insoluble fraction (Fig. 4D).

**Fig. 4:**
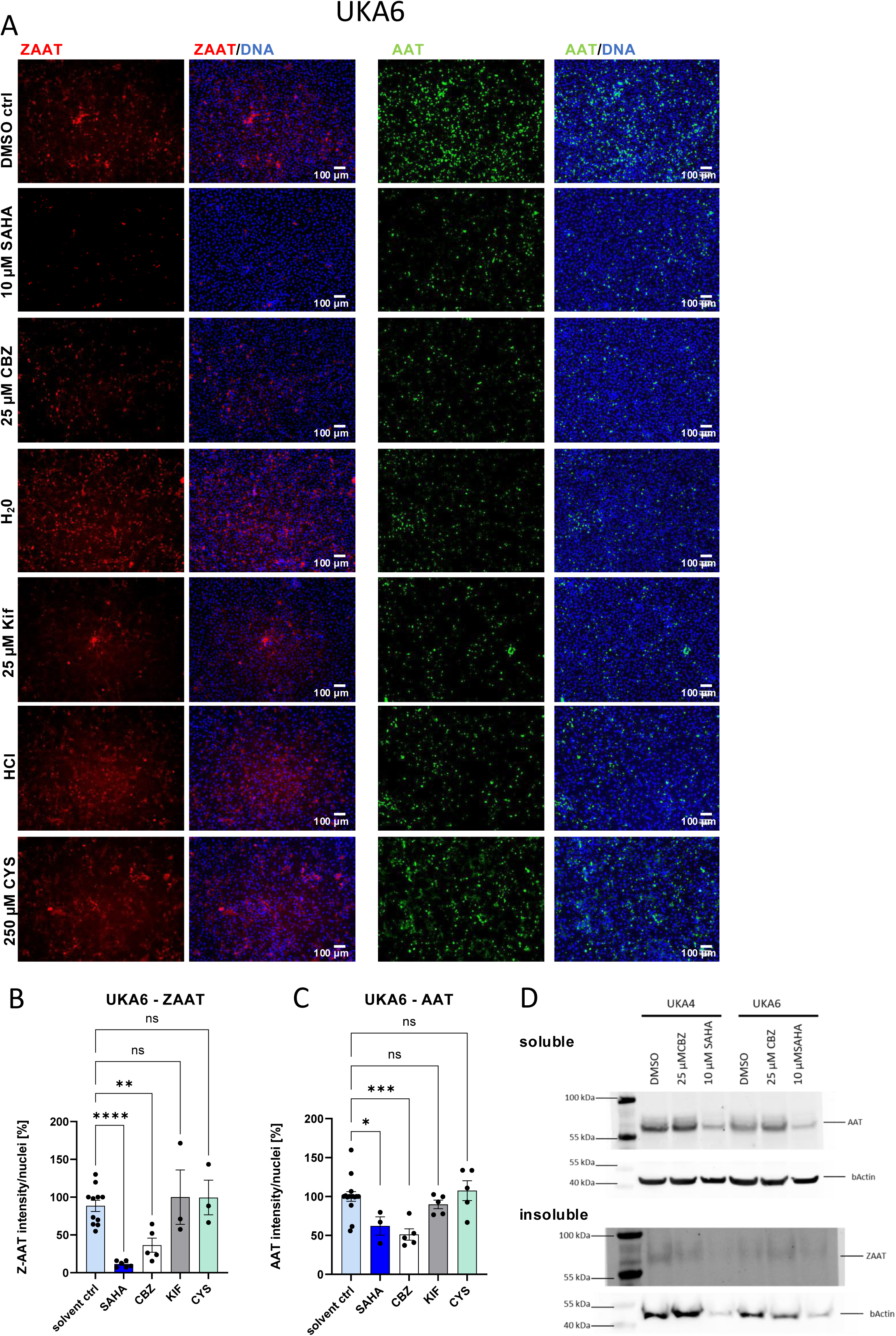
Small molecule treatment influences the amount of ZAAT and AAT in patient-derived HLCs. **(A)** ICC reveals changes in the amount of ZAAT (red) and AAT (green) after treatment with the indicated small molecules in relation to their respective controls (SAHA, CBZ: ctrl = DMSO, Kif: ctrl = H_2_O, CYS: ctrl = HCl). **(B,C)** Quantification of ZAAT (B) and AAT **(C)**, N= 3-11, bars represent mean ±SEM. Two-way ANOVA was performed to calculate significances (*p < 0.05, **p < 0.01, ***p < 0.001, ****p < 0.0001). **(D)** Western blot for the Triton X soluble (up) and insoluble (down) fraction of patient-derived HLCs indicate lower levels of AAT and ZAAT after SAHA treatment. bActin was used as housekeeping gene. n=1 (pool of 2 independent wells).

### SAHA treatment increases expression of HSP70 members and reduces ZAAT polymers

Based on the protein data, we decided to further study the effects of 25 µM CBZ and 10 µM SAHA on both patient-derived HLCs by bulk next generation sequencing. We identified 375 genes being exclusively expressed after CBZ and 1,500 after SAHA treatment. 493 and 577 genes were exclusively expressed in the DMSO control of the CBZ and SAHA treated cells, respectively (Suppl. Table S5A, Fig. 5A). Combining the exclusively and up-regulated genes after CBZ treatment, we identified 290 genes being significantly up-regulated and 233 genes being significantly down-regulated. Similarly, 245 genes were significantly up-, and 160 genes significantly down-regulated after SAHA treatment.

**Fig. 5:**
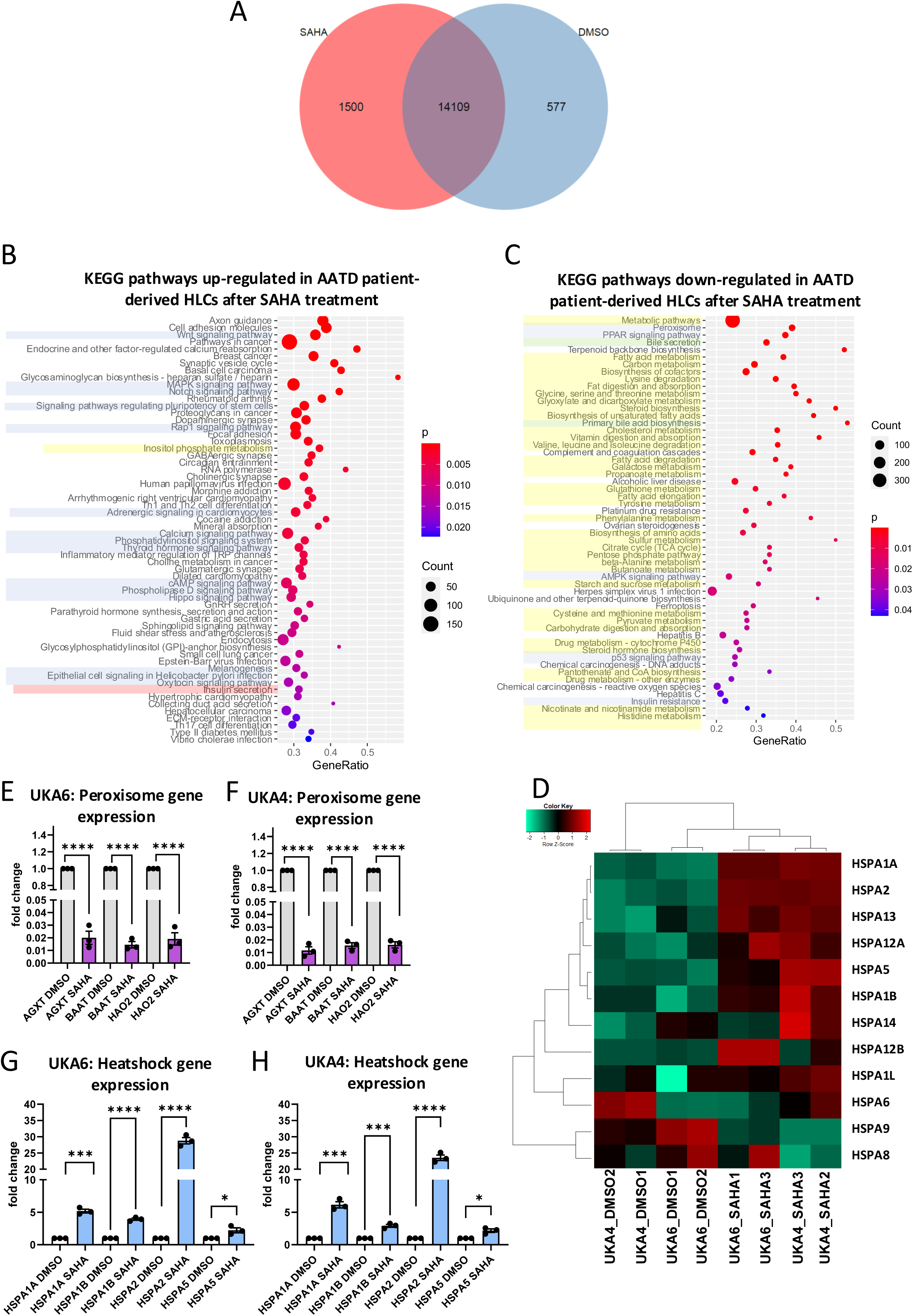
SAHA treatment increases expression of heat-shock genes and reduces expression of peroxisome genes. **(A)** Venn diagram indicating number of genes uniquely expressed in HLCs after SAHA treatment (1,500), uniquely expressed in control conditions (577) and commonly expressed (14,109) **(B)** KEGG pathways significantly up-regulated after SAHA treatment. **(C)** KEGG pathways significantly down-regulated after SAHA treatment. Pathways associated with signalling are highlighted in blue, with metabolism in yellow, with (bile) secretion in green, and with diabetes in red. For full data set, please refer to Suppl. Table S5. **(D)** Heatmap of heat-shock protein family 70 genes shows a general trend of up-regulation in HLCs after SAHA treatment. **(E-H)** Relative mRNA expression of heat shock protein genes (E,F) and peroxisome pathway genes (G,H) normalized to the housekeeping gene *RPLP0*. Bar plots show mean of triplicates ±SEM. Two-tailed Student’s t-test was performed to calculate significances (*p < 0.05, ***p < 0.001, ****p < 0.0001).

For both treatments, KEGG pathways analysis revealed up-regulation of several signalling pathways and down-regulation of metabolism-associated pathways (Fig. 5B,C, Suppl. Fig. S5B,C). Importantly, the latter comprised “Peroxisome” and “PPAR-signalling” for both treatments (Fig. 5C, Suppl. Fig. S5C), which are among the most strongly correlated pathways with ZAAT load in patient livers (Rosenberger et al., 2025). QPCR confirmed the downregulation of selected Peroxisome related genes after SAHA treatment (Fig. 5D,E) but not after CBZ treatment (Suppl. Fig. S5D,E).

As SAHA is known to enhance expression of heat-shock proteins (HSPs) (Di et al., 2013; Kuta et al., 2020) and we observed previously an interaction between GRP78 (encoded by *HSPA5*) and ZAAT polymers (Spivak *et al*., 2025), we looked deeper into the influence of SAHA on these genes in our HLCs. Importantly, we observed a consistent up-regulation of HSP70 family members after treatment of both patient-derived HLCs with SAHA (Fig. 5D, Suppl. Fig. S6) which was confirmed by qPCR (Fig. 5G,H). However, while we could detect GRP78 protein in the soluble and insoluble fraction of the patient-derived HLCs, we did not observe any quantitative changes in the protein level in association with SAHA or CBZ treatment (Suppl. Fig. S7). As the activity of GRP78 is heavily controlled by posttranslational modifications (Nitika et al., 2020), there might still be distinct levels of activity which we need to investigate deeper in the future.

## Discussion

In this study, we differentiated iPSCs from AATD patients into HLCs to test the potential of the four distinct compounds-CBZ, SAHA, KIF, and CYS- to reduce the levels of ZAAT polymer formation. These drugs have shown beneficial effects on ZAAT polymers in previous studies (Choi *et al*., 2013; Hidvegi *et al*., 2010; Karatas *et al*., 2021; Marcus and Perlmutter, 2000; Wilson *et al*., 2015) and except for KIF, they are all FDA approved for other indications (Grant et al., 2007; Sanchez et al., 2024; Santoro et al., 2025).

In line with previous studies, we saw that the AATD patient-derived iPSCs differentiated with similar efficiency towards HLCs as control cells (Choi *et al*., 2013; Kaserman *et al*., 2020; Kaserman *et al*., 2022; Kaserman and Wilson, 2018; Rashid *et al*., 2010; Segeritz *et al*., 2018; Tafaleng *et al*., 2015; Werder *et al*., 2021; Wilson *et al*., 2015; Yusa *et al*., 2011). While CYP3A4 activity was comparable between all 3 lines and could be significantly enhanced by induction with 10 µM rifampicin, CYP2D6 activity differed considerably between the lines. As CYP2D6 is known for its various genotypes that result in considerable variability in activity (Marez et al., 1997; Yang et al., 2017), it is very likely that UKA6 cells have a genotype associated with higher activity and did not differentiate more efficiently than the other cell lines, especially as the UM51 cells are known to have a rather weak intermediate metaboliser genotype (Bohndorf *et al*., 2017). In accordance with other studies, we observed higher levels of AAT in the patient-derived HLCs than in the control cells, (Wilson *et al*., 2015). Importantly, the pattern of AAT staining in the ICC differed considerably between patient-derived and control cells. While AAT is distributed homogeneously in the control cells, which is the typical pattern seen in healthy hepatocytes, it formed rather focused structures in the AATD-HLCs. In fact, these structures resembled those seen for the ZAAT polymers in these cells, which were undetectable in the control HLCs. Thus, the staining pattern already indicates polymer formation in the ZAAT HLCs.

To reveal more differences between patient and control HLCs, we analysed the transcriptome via NGS. In accordance with studies by Kaserman et al., we found “organelle” or “membrane-enclosed lumen” among the most highly up-regulated GO-CC terms in the patient-derived HLCs (Kaserman *et al*., 2022). This indicates aberrations in the intracellular transport system, which is a hallmark in AATD patients. This observation was supported by a high number of Golgi- and ER-related GO-CC terms being up-regulated in ZAAT-HLCs while many mitochondria-related GO-CC terms indicate a high energy demand of these cells which might be related to cell stress. When comparing KEGG-pathways between AATD- and control derived HLCs, we observed a general dysregulation of metabolism-associated pathways accompanied with a down-regulation of signalling associated pathways. Most importantly, we observed a down-regulation of secretion associated pathways, in particular “bile secretion” in the ZAAT HLCs, which might be related to the cholestatic phenotype seen in paediatric AATD-patients.

Among the 4 compounds that were supposed to be capable of influencing AATD polymers that we tested, only SAHA and CBZ significantly reduced ZAAT levels in both patient-derived HLCs. Interestingly, both compounds also reduced AAT-levels in these cells, indicating a feedback loop on AAT synthesis. This effect was particularly striking in the SAHA treated cells, where we saw a general reduction in the insoluble fraction containing the ZAAT-polymers, which even affected the housekeeping gene. This might be related to SAHA’s well-known capability to activate HSPs (Di *et al*., 2013; Kuta *et al*., 2020) which help to reduce the number of miss-folded proteins in general, thus reducing the insoluble fraction. While we previously demonstrated direct interactions of GRP78 with the ZAAT polymers in AATD-derived primary hepatocytes (Spivak *et al*., 2025), and saw an up-regulation of the corresponding gene *HSPA5* after SAHA-treatment in this study, we could not detect differences in GRP78 protein levels after SAHA treatment. This lack of correlation might be related to the fact that the activity of GRP78 is highly regulated by posttranslational modifications (Nitika *et al*., 2020) which requires further in-depth assessments in a follow-up study. Nevertheless, the general up-regulation of *HSP* genes might support the correct folding of mutated AAT, thus reducing cellular stress and increasing the probability of AAT secretion. This question warrants further studies directly assessing stress-associated pathways and measuring the amount of secreted AAT.

Besides influencing HSP-gene expression, we also observed that SAHA was capable of significantly down-regulating genes associated with the peroxisome proliferator pathway- a pathway which we previously identified as one of the most promising druggable pathways in ZAAT-patients (Spivak *et al*., 2025). Also here, further experiments are necessary to disentangle the role of the peroxisome proliferator pathway in AATD and its suitability as a drug target.

Overall, our data confirm that HLCs derived from iPSCs of ZAAT patients are a suitable model for AATD and can be used for the evaluation of potential novel drugs. In particular, our data indicate that SAHA might have beneficial effects for AATD-patients as it induced down-regulated expression of peroxisome proliferator-associated genes and up-regulated a host of HSPs such as *HSPA5*.

## Materials and Methods

### iPSC lines and cultivation

The use of two human iPSC lines from AATD patients (UKA4, UKA6 (Ncube *et al*., 2023)) and a control line (UM51 (Bohndorf *et al*., 2017)) was approved by the Ethical Committee of the medical faculty of Heinrich Heine University, Düsseldorf, Germany - approval numbers 2021-1627 and 5704 - respectively (Tabl. 1). iPSCs were maintained and expanded on Matrigel (Corning)-coated wells in StemMACS medium (Miltenyi) with penicillin/streptomycin (P/S) (Gibco) at 37°C, 5% O_2_ and 5% CO_2_ and daily medium changes. Cells were passaged 1:6-1:10 in colonies upon reaching approx. 70 % confluency using PBS w/o Ca^2+^, Mg^2+^.

### HLC differentiation

HLC differentiation was performed following our optimized three-step protocol (Loerch *et al*., 2024). In brief, 1.04 × 10^5^ iPSCs/cm^2^ were seeded as single cells in StemMACS with 10 μM Y27632 (Tocris) and incubated at 37°C, 5% CO_2_. 24 h after seeding, StemMACS was removed and definitive endoderm (DE) medium consisting of RPMI 1640, 2% B27 w/o retinoic acid, 1% Glutamax (Glx) (all Gibco), 2 µM doxycycline (Doxy) (Fisher Scientific), and 100 ng/ml Activin A (Stem Cell Technologies) was added for 3 days with daily medium change. For the first day, 2.5 μM Chir99021 (Stemgent) was included. On day 4 of differentiation, medium was changed to hepatic endoderm (HE) medium (DMEM/F12, 20% Knockout Serum Replacement (KO-SR, 0.01% β-mercaptoethanol (all Gibco), 1% DMSO (Corning)). Medium was changed daily. From day 8 on, cells were fed every other day with HLC medium (DMEM/F12, 8% FBS, 8% tryptose phosphate broth, 1% Glx (all Gibco), 2 µM Doxy, 1 μM Insulin (Sigma-Aldrich), 10 ng/ml HGF (Peprotech), 25 ng/ml Dexamethasone (Sigma-Aldrich), 20 ng/ml oncostatin M (Immunotools) and 20 μM Forskolin (Tocris).

HLCs generated from AATD-patient iPSC were treated with CBZ (Sigma), SAHA (Cayman), KIF (Cayman), CYS (Sigma) for 48 h, starting on day 19 of differentiation. HLCs were incubated in HLC medium containing all supplements and the indicated concentrations of the four factors or the appropriate solvent control at a dilution of 1:1000. DMSO was used as solvent control for CBZ and SAHA, H_2_O for KIF and HCl for CYS.

### Immunocytochemistry (ICC)

For Immunocytochemistry, cells were fixed with 4% PFA and permeabilized for 10 min with 0.5% Triton-X-100 (Sigma-Aldrich) in PBS. After blocking with 3% BSA/PBS, primary antibodies (Suppl. Table S1) were applied overnight at 4°C. Cells were washed 3 times and incubated with fluorescence labelled secondary antibody (Suppl. Table S2) against the respective host species IgG (Life technologies) for 2 h at RT. Hoechst 33258 was used to stain the nuclei. Pictures were taken with a LSM 700 microscope (Zeiss) and processed with ZENsoftware (Zeiss). Olympus cellSens Dimension Desktop 1.16 software was used for quantification from ICC pictures and significances were calculated with two-way ANOVA, *p < 0.05, **p < 0.01; ***p < 0.001, ****p < 0.0001.

### Real time quantitative PCR

RNA isolation was performed using the Direct-zol RNA MiniPrep (ZYMO Research) according to the manufacturer’s instructions. 500 ng of RNA was reverse transcribed to cDNA using the TaqMan Reverse Transcription Kit (Applied biosystems). PowerTrack SYBR Green Master Mix (Applied biosystems) was used for qPCR. For each gene, 5 ng of cDNA were amplified in technical triplicates using a VIIA7 machine (Lifetechnologies). All primers were ordered from Eurofins (Suppl. Table S3). Relative mRNA expression was calculated as the log2-fold change between the gene of interest and the housekeeping gene *RPLP0*. Significances were calculated with two-tailed unpaired Student’s t-tests.

### Cytochrome P450 activity assay

CYP activity assays were performed with the respective P450-Glo kits from Promega. On day 20 of differentiation, HLCs were incubated w/ and w/o 10 μM Rifampicin to induce CYP3A4 activity. On day 21, William’s E medium (Gibco) containing 3 μM Luciferin-IPA (CYP3A4) or 30 μM of Luciferin-ME EGE was added to the cells. HLCs were incubated for 1 hour at 37 °C to allow the CYP-mediated conversion of the respective substrates from pro-luciferin to luciferin. Medium was harvested and mixed 1:1 with the detection reagent containing luciferase. After 20 min incubation at RT in the dark, luminescence was measured using the Lumat LB 950 and subsequently normalized to protein content.

### Indocyanine green dye (ICG) uptake and release

HLCs were incubated for 30 min at 37 °C with HLC medium containing 0.5 mg/ml ICG. After incubation, medium containing ICG was removed and cells were washed 3x with HBSS. Brightfield pictures were taken before fresh HLC medium with all supplements was added. Cells were incubated for 6 h at 37 °C to allow ICG release. Medium was removed and cells were washed with HBSS before brightfield pictures were taken.

### Western Blot

For standard Western Blot (Fig. 2B), cells were lysed with RIPA buffer (Sigma-Aldrich) containing protease and phosphatase inhibitors (both Roche). Protein concentration was determined using the BCA Protein Assay Kit (Pierce). 20 µg of protein was loaded on a 10% acrylamide (Roth) gel. To detect ZAAT (Fig. 4D, Suppl. Fig. S7), cells were lysed in PBS containing 1 % Triton-X-100 (Roth), 5 mM EDTA (Roth) and proteinase and phosphatase inhibitors. The insoluble and soluble fractions were separated by centrifugation at 20,000 *x g*, at 4 °C for 15 min. Soluble fractions were denatured in Laemmli buffer and cooked at 95 °C for 5 min prior to use. The insoluble fractions were homogenized by vortexing with glass pearls in 4x Laemmli buffer for 2 h at 4°C and cooked at 95 °C for 10 min. Proteins were separated and transferred onto 0.45 µm nitrocellulose membranes (Amersham) by wet blot. Successful transfer was confirmed by Ponceau (Fisher Scientific) staining. Membranes were blocked in TBS with 1% Tween 20 (TBST) containing 5% milk powder for 1 – 2 h at RT. Primary ABs were diluted according to Suppl. Table S1 in blocking buffer and membranes were incubated overnight at 4 °C. Membranes were washed 3x with TBS-T and incubated with fluorophore-coupled secondary ABs diluted in blocking buffer according to Suppl. Table S2 for 2 h at RT. Afterwards, membranes were washed 3x with TBS-T 1X for 5 min. For HRP detection, ECL Western Blotting Substrate (Pierce) containing HRP substrate solution was used and chemiluminescence signals were detected using the Fusion FX (Vilber Lourmat). To detect fluorophore-coupled secondary antibodies the fluorescence signals were detected at the Odyssey CLx (LI-COR Biosciences) or the ChemiDoc MP Imaging system (Bio-Rad).

### Next generation sequencing

3’RNA-Seq was performed on a NextSeq2000 sequencing system (Illumina) at the core facility Biomedizinisches Forschungszentrum Genomics and Transcriptomics laboratory (BMFZ-GTL) of Heinrich-Heine University Duesseldorf. Significant differential expression was defined by a p-value < 0.05 from the limma test and a fold change > 1.5 for up-regulated genes or < 0.667 for down-regulated genes, taking into account genes expressed in at least one condition (Suppl. Tables S4 and S5). Genes with more than 5 reads were defined as expressed. For details, please refer to supplementary Materials and Methods. NGS data for the control sample have been published in (Loerch et al., 2026) (sample Cntrl 1, DMSO treated). Data will be made available via the GEO server upon acceptance of the manuscript.

### GO and pathway analysis

Subsets of genes expressed exclusively in one condition in the Venn diagram analysis and up- and down-regulated genes according to the criteria for differentially expressed genes mentioned above were subjected to over-representation analysis of gene ontologies (GOs) and KEGG (Kyoto Encyclopedia of Genes and Genomes) pathways (Kanehisa et al., 2017). The hypergeometric test built-in in the R base package was used for over-representation analysis of KEGG pathways, which had been downloaded from the KEGG database in February 2023. The GOstats R package (Falcon and Gentleman, 2007) was employed to determine over-represented GO terms. The most significant GO terms and KEGG pathways were displayed in dotplots via the R package ggplot2 (Wickham, 2009).

## Resource availability

NGS data will be made available via the GEO server upon acceptance of the manuscript.

## Acknowledgments

J.A. acknowledges funding from the Medical Faculty of Heinrich-Heine University Düsseldorf, C.L. and N.G. were funded by the Else Kroener-Fresenius Foundation – 2020_EKEA.64. N.G. acknowledges funding from the Ministry for Culture and Sciences North Rhine Westfalia – 005-2305-0038. The graphical abstract was created in BioRender. Graffmann, N. (2026) https://BioRender.com/4a9lq5q

## Author contributions

**N.G.:** Conceptualization, Data curation, Formal analysis, Funding acquisition, Investigation, Methodology, Validation, Visualization, Writing – Original draft, Writing, review & editing, **R.H.:** Data curation, Formal analysis, Investigation, Methodology, Validation, Visualization, Writing – Original draft, Writing, review & editing, **C.L.:** Data curation, Formal analysis, Investigation, Methodology, Validation, Visualization, Writing – Original draft, Writing, review & editing, **M.F.:** Resources, Validation, Writing – Original draft, Writing, review & editing, **W.W.:** Data curation, Formal analysis, Investigation, Validation, Visualization, Writing – Original draft, Writing, review & editing, **P.S.:** Conceptualization, Funding acquisition, Resources, Supervision, Validation, Writing – Original draft, Writing, review & editing, **J.A.:** Conceptualization, Funding acquisition, Supervision, Validation, Writing – Original draft, Writing, review & editing,

## Abbreviations

AAT: Alpha-1-Antitrypsin
AATD: Alpha-1-Antitrypsin deficiency
AFP: Alpha-Fetoprotein
ALB: Albumin
CC: Cellular compartment
CBZ: Carbamazepine
COPD: Chronic obstructive pulmonary disease CYP Cytochrome P450
CYS: Cysteamine
DE: Definitive endoderm
Dex: Dexamethasone
ECAD: E Cadherin
ER: Endoplasmic reticulum
FDA: Food and drug administration (U.S.)
GOs: Gene ontologies
GRP78: 78 kDa glucose-regulated protein
HCC: Hepatocellular carcinoma
HDACi: Histone deacetylase inhibitor
HE: Hepatic endoderm
HGF: Hepatocyte growth factor
HLCs: Hepatocyte like cells
HNF4a: Hepatocyte nuclear factor 4alpha
HSP: Heat-shock protein
HSPA5: Heat-shock protein family A (Hsp70) member 5
ICC: Immunocytochemistry
iPSCs: Induced pluripotent stem cells
KEGG: Kyoto Encyclopedia of Genes and Genomes
KIF: Kifunensine
MAPK: Mitogen-activated protein kinase
NE: Neutrophile elastase
NGS: Next generation sequencing
OSM: Oncostatin M
PI: Protease inhibitor
PPAR: Peroxisome proliferator activated-receptor
P/S: Penicillin/Streptomycin
RAP1: Ras-proximate 1
SERPIN: Serin protease inhibitor
SNP: Single nucleotide polymorphism
STRING: Search Tool for the Retrieval of Interacting Genes/Proteins
qRT-PCR: Quantitative reverse transcription PCR
R.L.U.: Relative light units
RNA-seq: RNA sequencing
RPLP0: Ribosomal Protein Lateral Stalk Subunit P0
SAHA: Suberoylanilide hydroxamic acid
SOX: SRY-related HMG-box genes
UPR: Unfolded protein response
ZAAT: Z variant of AAT

## Supplemental information

**Document S1.** Figures S1–S7, Supplementary methods with Tables S1-S3

**Supplementary table S4: NGS data_HLC AATD vs ctrl**

**Supplementary table S5: NGS data_HLC AATD SAHA_CBZ**

