## Supplementary figures and Methods for "SAHA increases chaperone expression and reduces Z-alpha-1-antitrypsin polymers in a patient specific iPSC-based liver model for alpha-1-antitrypsin deficiency"

### Suppl. Fig. 1

**A** STRING network of genes common in at least 2 of the transcriptomic pathways down-regulated in AATD-patient derived HLCs

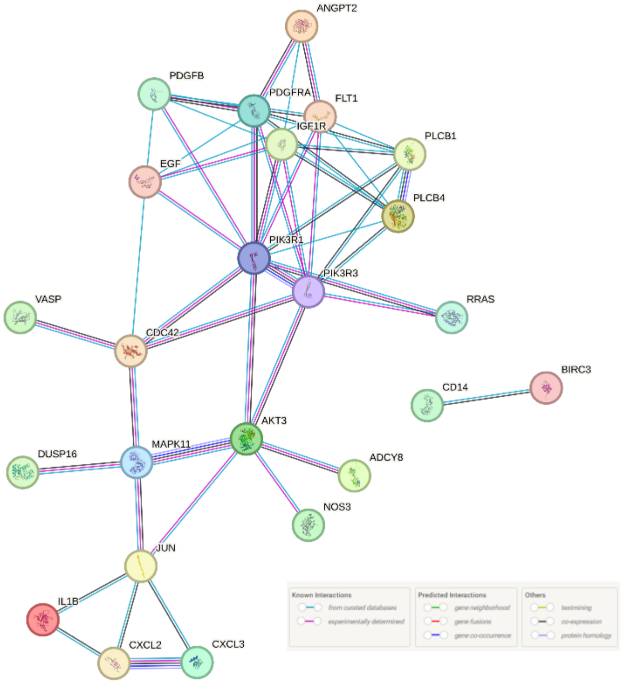


**Suppl. Fig. 1: STRING network of genes common in at least 2 of the transcriptomic pathways down-regulated in AATD patient-derived HLCs**

### Suppl. Fig. 2

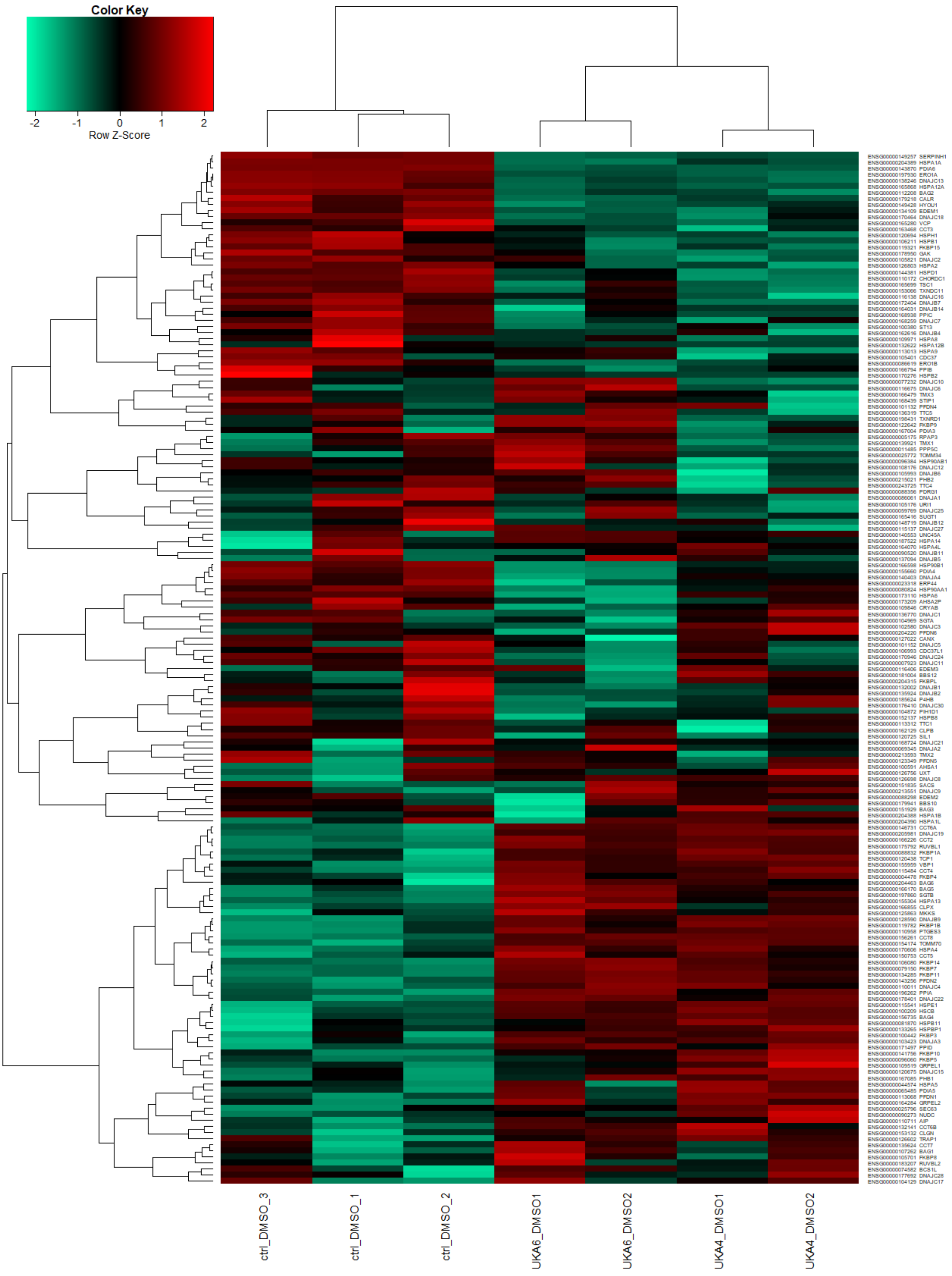


**Suppl. Fig. 2: Heatmap of heat-shock protein genes shows clear clustering according to AAT genotype**

Suppl Fig. 3 UKA 6

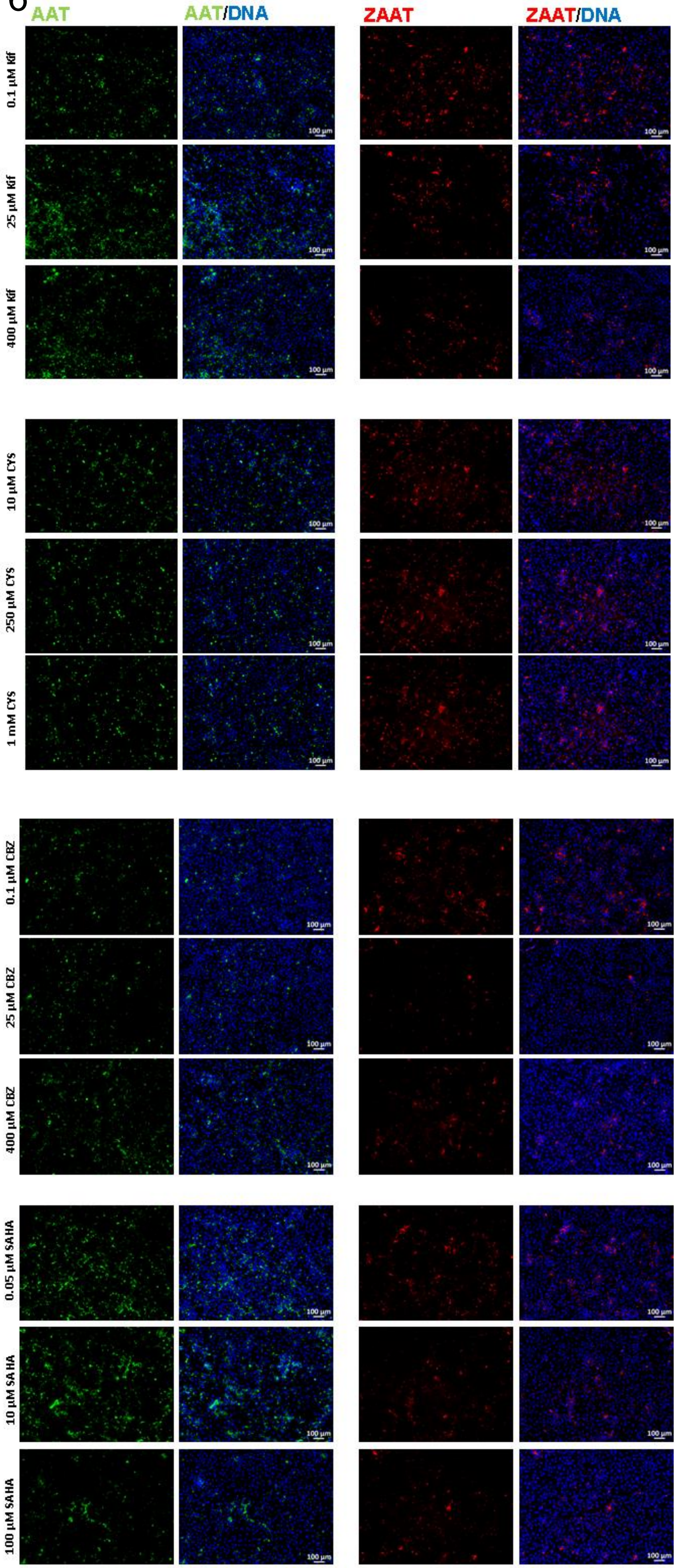

**Suppl. Fig. 3: Concentration testing of the 4 compounds.** ICC was performed to test changes in the amount of ZAAT (red) and AAT (green) after treatment with the indicated small molecules in relation to their respective controls (SAHA, CBZ: ctrl = DMSO, Kif: ctrl = $H_2O$, CYS: ctrl = HCl). For each compound in total 9 different concentrations ranging between 0.1 µM – 400 µM for CBZ and KIF, 0.05 µM – 100 µM for SAHA, and 10 µM – 1 mM for CYS were tested on UKA6 HLCs. Depicted are the lowest and highest concentrations as well as the concentration selected for the experiments.

### Suppl. Fig. 4 UKA4

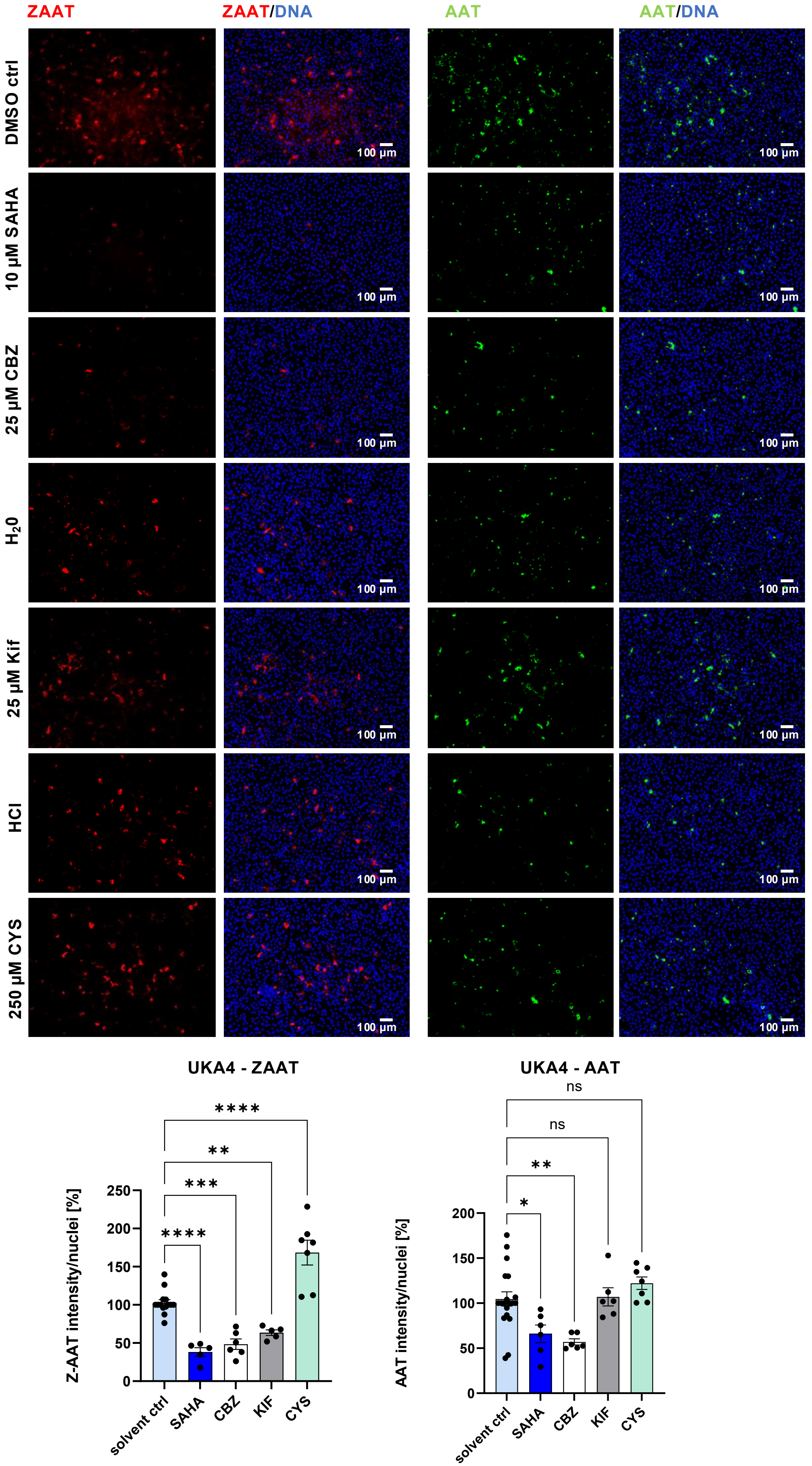

**Suppl. Fig. 4: Small molecule treatment influences the amount of ZAAT and AAT in patient-derived HLCs. (A)** ICC reveals changes in the amount of ZAAT (red) and AAT (green) after treatment with the indicated small molecules in relation to their respective controls (SAHA, CBZ: ctrl = DMSO, Kif: ctrl = $H_2O$, CYS: ctrl = HCl). **(B)** Quantification of ZAAT and AAT, N= 5-19, bars represent mean ±SEM. Two-way ANOVA was performed to calculate significances (*$p < 0.05$, **$p < 0.01$, ***$p < 0.001$, ****$p < 0.0001$).

Suppl. Fig. 5

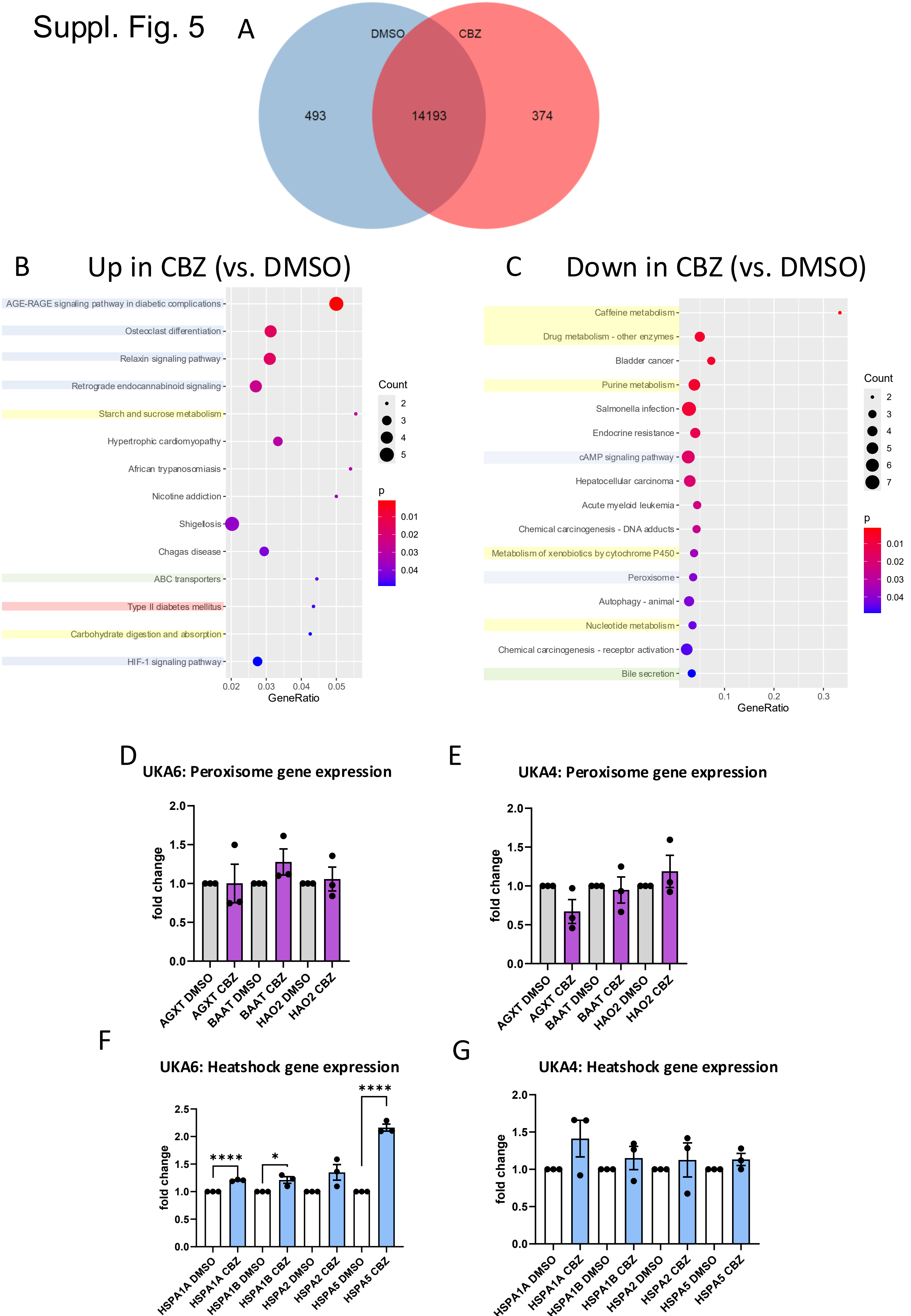
A
DMSO
CBZ
493
14193
374
B
Up in CBZ (vs. DMSO)
AGE-RAGE signaling pathway in diabetic complications
Osteoclast differentiation
Relaxin signaling pathway
Retrograde endocannabinoid signaling
Starch and sucrose metabolism
Hypertrophic cardiomyopathy
African trypanosomiasis
Nicotine addiction
Shigellosis
Chagas disease
ABC transporters
Type II diabetes mellitus
Carbohydrate digestion and absorption
HIF-1 signaling pathway
GeneRatio
Count
p
C
Down in CBZ (vs. DMSO)
Caffeine metabolism
Drug metabolism - other enzymes
Bladder cancer
Purine metabolism
Salmonella infection
Endocrine resistance
cAMP signaling pathway
Hepatocellular carcinoma
Acute myeloid leukemia
Chemical carcinogenesis - DNA adducts
Metabolism of xenobiotics by cytochrome P450
Peroxisome
Autophagy - animal
Nucleotide metabolism
Chemical carcinogenesis - receptor activation
Bile secretion
GeneRatio
Count
p
D
UKA6: Peroxisome gene expression
fold change
AGXT DMSO
AGXT CBZ
BAAT DMSO
BAAT CBZ
HAO2 DMSO
HAO2 CBZ
E
UKA4: Peroxisome gene expression
fold change
F
UKA6: Heatshock gene expression
fold change
HSPA1A DMSO
HSPA1A CBZ
HSPA1B DMSO
HSPA1B CBZ
HSPA2 DMSO
HSPA2 CBZ
HSPA5 DMSO
HSPA5 CBZ
G
UKA4: Heatshock gene expression
fold change

**Suppl. Fig. 5: CBZ treatment increases the expression of genes associated with signalling pathways and reduces metabolism associated genes. (A)** Venn diagram indicating number of genes exclusively expressed in HLCs after CBZ treatment (347). uniquely expressed in control conditions (493) and commonly expressed (14193) **(B)** KEGG pathways significantly up-regulated after CBZ treatment. **(C)** KEGG pathways significantly down-regulated after CBZ treatment. Pathways associated with signalling are highlighted in blue, with metabolism in yellow, with (bile) secretion in green, and with diabetes in red. For full data set, please refer to Suppl. Table 5. **(D-G)** Relative mRNA expression of heat shock protein genes (D,E) and peroxisome pathway genes (F,G) normalized to the housekeeping gene *RPLP0*. Bar plots show mean of triplicates ±SEM. Two-tailed Student's t-test was performed to calculate significances (*p < 0.05, ****p < 0.0001).

### Suppl. Fig. 6

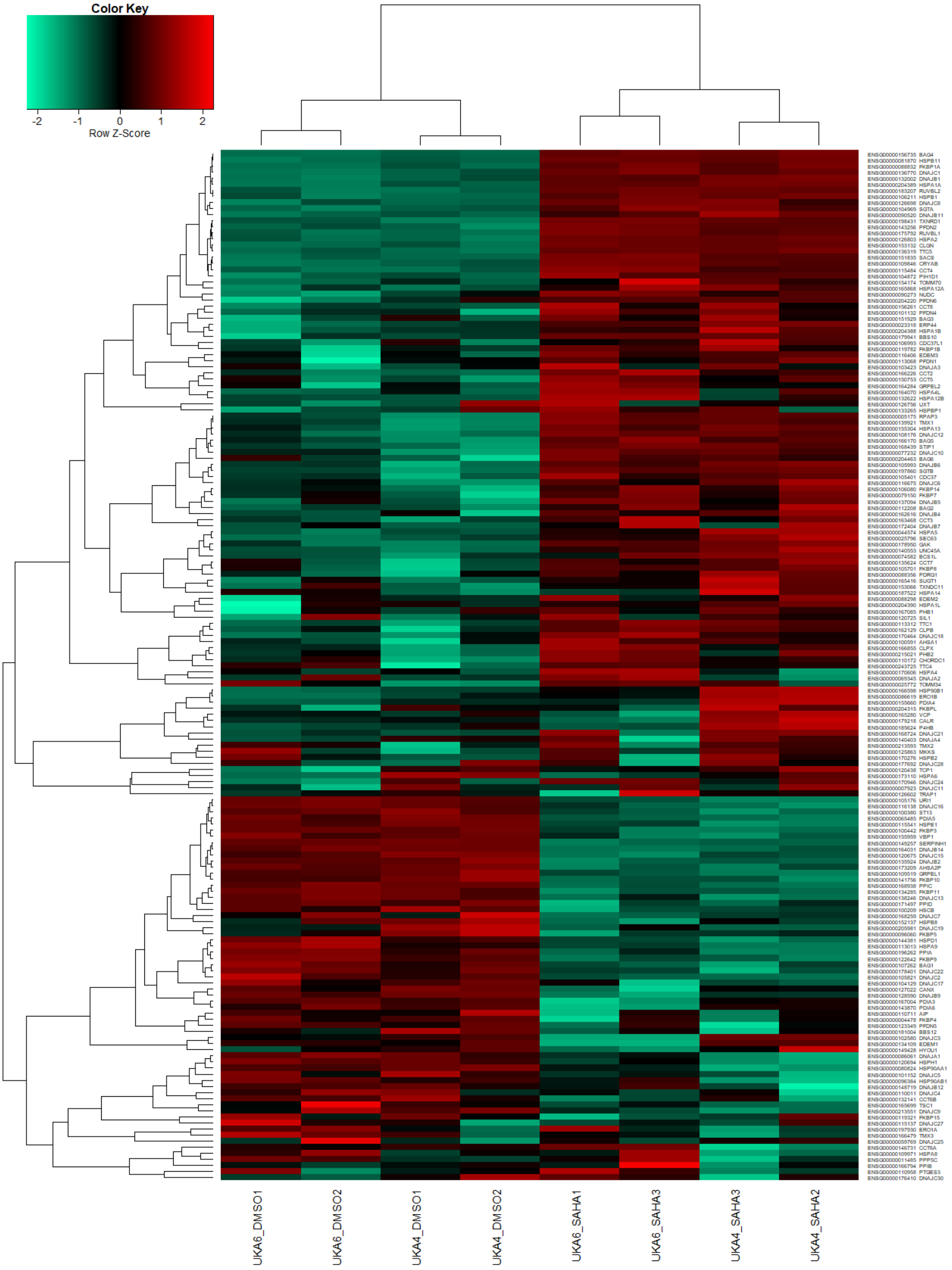


**Suppl. Fig. 6: Heatmap of heat-shock protein genes shows clear clustering according to SAHA vs DMSO treatment**

### Suppl. Fig. 7

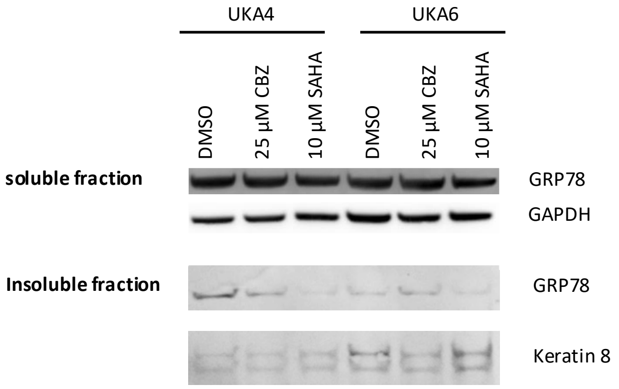


**Suppl. Fig. 7:** Western blot for the Triton X soluble (up) and insoluble (down) fraction of patient-derived HLCs indicate similar levels of GRP78 after CBZ or SAHA treatment. GAPDH and keratin 8 served as respective housekeeping genes. n=1 (pool of 2 independent wells).

#### Supplementary Materials and Methods

**Supplementary Table 1**: List of primary antibodies used for ICC and WB

| Primary Antibody | Species | Dilution ICC | Dilution WB | Company |
|---|---|---|---|---|
| **anti-AAT** | goat | 1:1,000 | 1:3,000 | GeneTex |
| **anti-β-Actin** | mouse | | 1:5,000 | Cell Signalling Technology |
| **anti-ALB** | mouse | 1:100 | 1:5,000 | Sigma |
| **anti-GAPDH** | rabbit | 1:200 | 1:1,000 | Cell Signalling Technology |
| **anti-HNF4α** | rabbit | 1:250 | | Abcam |
| **anti-GRP78** | mouse | | 1:1,000 | BD |
| **anti-Keratin 8** | mouse | | 1:1,000 | Progen |
| **anti-γ-Tubulin** | mouse | | 1:1,000 | Sigma |
| **anti-ZAAT** | mouse | 1:50 | 1:50 – 1:500 | HycultBiotech |

**Supplementary Table 2**: List of secondary antibodies used for ICC and WB

| Secondary Antibody | Species | Dilution | Company |
|---|---|---|---|
| **anti-mouse 488** | goat | 1:500 | Invitrogen |
| **anti-mouse 555** | goat | 1:500 | Invitrogen |
| **anti-rabbit 488** | goat | 1:500 | Invitrogen |
| **anti-rabbit 647** | goat | 1:500 | Invitrogen |
| **anti-goat 488** | donkey | 1:500 | Invitrogen |
| **anti-goat 680RD** | donkey | 1:10,000 | LI-COR Biosciences |
| **anti-mouse 680RD** | donkey | 1:10,000 | LI-COR Biosciences |

**Supplementary Table 3**: List of primers

| Primer | | Sequence 5'→ 3' | Product size (bp) |
|---|---|---|---|
| **AAT** | forward | GGTCACAGAGGAGGCACCC | 76 |
| | reverse | AGTCCCTTTCTCGTCGATGGT | |
| ***ALB*** | forward | AGCTGTTATGGATGATTTCGCAG | 77 |
| | reverse | CCTCGGCAAAGCAGGTCTC | |
| ***AFP*** | forward | AGCAGCTTGGTGGTGGATGA | 88 |
| | reverse | CCTGAGCTTGGCACAGATCCT | |
| ***AGXT*** | forward | ACCATCACACAATCCCCGTC | 222 |
| | reverse | CTCTCCAGTCATAGCCAGCG | |
| ***BAAT*** | forward | CCTCCTTGGCCTTGGCTTAC | 201 |
| | reverse | TGGCTGTGACTTGCTTTAGGT | |
| ***CYP3A4*** | forward | GTGACTTTGCCCATTGTTTAGAAAG | 79 |
| | reverse | CAGGCGTGAGCCACTGTG | |
| ***CYP3A7*** | forward | GATTCTGTACGTGCATTGTGCTC | 77 |
| | reverse | ATTTGGTCATCTCCTCTATATTACCAAGT | |
| ***HAO2*** | forward | CTGCAAGGGTGAACATGGTG | 106 |
| | reverse | TCGATTGATCTCAGCGACCG | |
| ***HNF4α*** | forward | GCACTCGAAGGTCAAGCTA | 157 |
| | reverse | GACTCACACACATCTGCGA | |
| ***HSPA1A*** | forward | AGCTGGAGCAGGTGTGTAAC | 154 |
| | reverse | CAGCAATCTTGGAAAGGCCC | |
| ***HSPA1B*** | forward | GCAGGTGTGTAACCCCATCA | 177 |
| | reverse | GAGTCCCAACAGTCCACCTC | |
| ***HSPA2*** | forward | GACGACATTGACCGGATGGT | 154 |
| | reverse | CGCTAATCTTGCCCCTCAGT | |
| ***HSPA5*** | forward | CCGAGGAGGAGGACAAGAAGG | 229 |
| | reverse | TCAAAGACCGTGTTCTCGGG | |

| | | | |
|---|---|---|---|
| ***RPLP0*** | forward | TCGACAATGGCAGCATCTAC | 110 |
| | reverse | ATCCGTCTCCACAGACAAGG | |

**Next generation sequencing (NGS)**

The RNA-seq raw data in FASTQ format received from the BMFZ-GTL core facility was aligned against the GRCh38 genome with the HISAT2 (version 2.1.0) alignment software[1]. The essentials of this procedure were described in our previous publication[2] with marginal changes in the annotation files and in the HISAT2 command, adapted from the parameter optimizations of Barruzzo et al. [3]:

hisat2 -p 7 -N 1 -L 20 -i S,1,0.5 -D 25 -R 5 --mp 1,0 --sp 3,0 -x /home/ww/hisatindex/grch38_r109 -U input.fastq.gz -S output.sam

Sorting was achieved by the SAMtools software [4] and summarization per gene by the subread (1.6.1) featureCounts software [5] against the annotations from Ensembl Homo_sapiens.GRCh38.109.gtf using parameters –t exon –g gene_id. For filtering expressed genes a threshold of CPM (counts per million) > 1 in at least one sample was applied before limma voom normalization [6,7] in R/Bioconductor [8].
